# Inuit Hunt as a Platform for Observing Narwhals (*Monodon monoceros*) in Inglefield Bredning (Kangerlussuaq), Greenland

**DOI:** 10.1101/2025.09.03.673629

**Authors:** Monica Ogawa, Evgeny A. Podolskiy

## Abstract

Direct observations of narwhals are scarce but needed for understanding this ecologically and culturally important species. Here, we describe the first boat-based observations of narwhals in their key summering ground in Greenland (Inglefield Bredning), collected during an Inuit hunt and enhanced by drone air support. During 3—8 August 2024, 506 narwhal observations were made from a semi-stationary boat at the head of the fjord, of which 58 were filmed with a drone. Boat observations near the north side of the fjord indicated that narwhals preferentially traveled outward to the west, with a clear link to currents. The presence of narwhals was more likely in the second half of a day but was highly intermittent, with waiting times for observers reaching as much as 15–28 h between sightings, highlighting the patience needed to observe and catch a narwhal. We also recorded a motionless sleep-like behavior at the surface, known as *pugginnartoq* (Greenlandic) for narwhals. Aerial drone support was useful for revealing unseen-from-sea-surface features and behaviors, which we describe as potentially interesting for future investigation. For example, drone imagery revealed that 71±7% of narwhals had tusks, with a mean tusk-to-body-length ratio of 0.23±0.01. Overall, this report shows that hunting expeditions integrated with scientific methods provide important insights and inspire further work.

## Introduction

The narwhal (*Monodon monoceros*), one of the most northerly distributed cetaceans in the world, is an ecologically, economically, and culturally important marine mammal in the Arctic region, including Greenland or Kalaallit Nunaat (*Heide-Jørgensen, 2024*). Inglefield Bredning Fjord (Kangerlussuaq) in northwest Greenland is ~100 km long and is a key summering ground for narwhals, with an estimated 2,874 individuals (cv = 0.21; 95% Cl = 1,938–4,354; 2019) coming to the fjord in summer (*Hansen et al., 2024*). Nevertheless, due to the skittish and shy nature of the local narwhal subpopulation, direct observations of their presence and behavior remain challenging. In other regions, like Canada and East Greenland, observations could be made visually from the coast (e.g., *Marcoux et al., 2009*) or via long-term biologging (e.g., *Podolskiy* and *Heide-Jørgensen, 2022*). However, to date, the observations of living narwhals in Inglefield Bredning have been limited to indirect approaches such as two short passive acoustic campaigns (*Miller et al., 1995; Podolskiy and Sugiyama, 2020*), abundance snapshots from aerial surveys (*Born et al., 1994; Heide-Jørgensen et al., 2010*; *Hansen et al., 2024*), short-term biologging (*Heide-Jørgensen and Laidre, 2006*), and ongoing autonomous long-term passive acoustic monitoring (e.g., *Podolskiy et al., 2022*).

In addition to the general knowledge gap about the presence and behavior of narwhals in their summering ground, there is a major disagreement between scientists and hunters about the abundance of narwhals in Greenland (*Heide-Jørgensen, 2024*). The two parties disagree about the status of the stocks (e.g., *Hansen et al., 2024*). To reconcile Indigenous knowledge and scientific methods, it is increasingly important to record hunters’ perspectives, especially since wildlife management requires both approaches. While standard boat-based visual surveys of narwhals have not been undertaken in Inglefield Bredning, the fjord is one of the last areas where traditional narwhal hunting is practiced using kayaks/qajaqs (also practiced in East Greenland’s Kangerlussuaq, *Heide-Jørgensen, 2024*). The hunting strategy is to wait next to an iceberg, sometimes for days, for an opportunity to quietly pursue a narwhal and harpoon it (Fig. 1). This narwhal hunt provides a rare opportunity to closely and directly observe the natural behavior of narwhals. While little is known of narwhal behavior, Indigenous people who have observed them in this fjord for generations have significant knowledge of them, which remains to be documented. The present study aims to show how much science can learn quantitatively and qualitatively by using an Inuit hunt as a marine observation platform. Our objectives are twofold: first, to demonstrate that the hunt can yield unique insights about some of Greenland’s least studied subpopulation of narwhals, and second, to show that aerial drone support helps to identify potentially interesting research questions without disturbing the hunt and narwhals. To enable a close approach to narwhals, visual sightings, and collection of video evidence, we collaborated with local hunters and joined their hunting efforts in August 2024. This fieldwork was part of a comprehensive study of narwhals, including stomach content analysis, long-term hydroacoustic monitoring, and interviews with hunters regarding marine mammals *(Ogawa, 2025; Podolskiy et al., 2022)*. Our field data yielded unique observations, adding to the scarce records of narwhal behavior. Considering the extreme sensitivity of narwhals to changing climate, human disturbance, and hunting pressure (*Heide-Jørgensen et al., 2020, 2021, 2024*), this is a timely contribution.

**Fig. 1.**
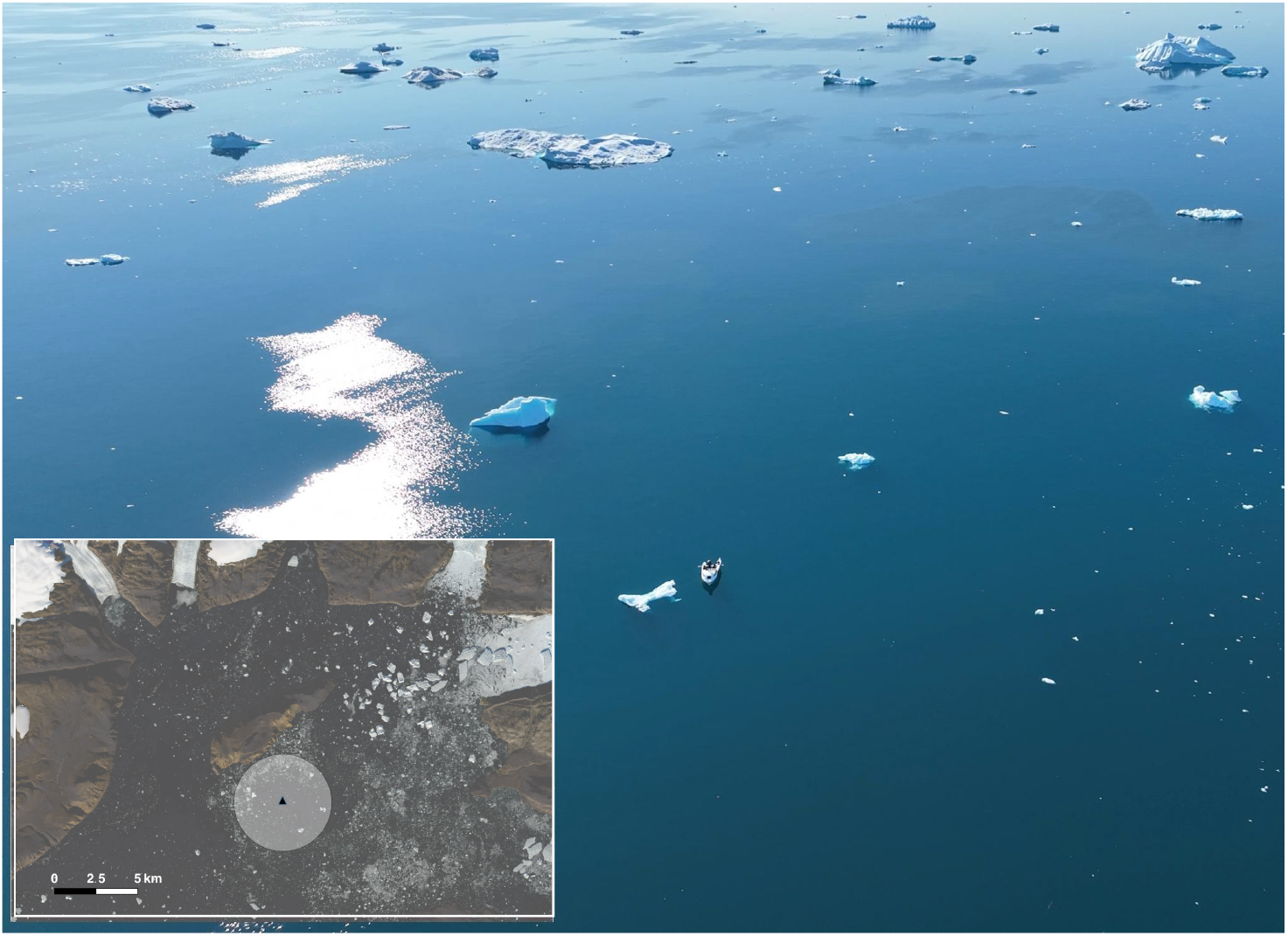
Narwhal observations at the head of Inglefield Bredning, Northwest Greenland (credits: M. Ogawa and T. Peary). The circle on the north-oriented satellite imagery (LANDSAT8, 7 September 2023) encompasses all the boat positions during the hunt

## Materials and methods

Visual observations were undertaken during a narwhal hunt on 3–8 August 2024, primarily between Qeqertaarsuusarsuaq (Josephine Peary Island) and the island of Qeqertat (also known as Harvard Island). While the boat shifted slightly with currents, all observations occurred within a 2.5 km radius of 77.5741°N, 66.7310°W (Fig. 1a). As this period coincided with the midnight sun, the observations were undertaken continuously (24 h per day), but had to be terminated on 9 August due to an approaching storm. During the observations, conditions were clear and calm, with five observers, including three experienced Inuit, constantly monitoring the sea surface with binoculars (×8). When narwhals were detected, the time, number of individuals, position (approximate distance and bearing from the front of the boat), behavior, detailed notes about boat orientation, and hunting activities were recorded. The distance between the observed narwhals and the boat was estimated visually. During sightings, some narwhal behavior was recorded opportunistically using a hand-held camera with a telescopic lens (Olympus E-M5 Mark III). Drone videos (4K, 3840 × 2160 px resolution) were recorded using a DJI Mavic 3 drone (UAV) with an 8.3 MP camera, from a mean altitude of 71 m (standard deviation, SD 12 m) with a variable angle to the sea surface (~70% of the videos were oblique). Mean coordinates of the drone videos were 77.5691°N, 66.6824° W. Considering potential interference with the hunt and the unknown sensitivity of narwhals to drone noise, the altitude was relatively high compared to other drone-based observations of cetaceans (e.g., 25 m for gray whales; *Torres et al., 2018*).

## Results and discussion

### Boat-based observations

#### Sighting statistics

In total, 506 narwhal observations were made from the boat between the evenings of 3 and 8 August (Fig. 2). Since it is difficult to confirm that the 506 narwhals observed were unique individuals, we cannot exclude that some narwhals were observed multiple times. The median pod size was five individuals. It is possible that the observers missed individuals beneath the surface of the water however, this number is consistent with a previous sighting effort undertaken in the 1980s from the coast near Qeqertat (*Born et al., 1994*), but larger than the expected group size of 3.3 estimated from aerial surveys in 2019 (*Hansen et al., 2024*) and the average cluster size of 3.5 seen from the coast in Koluktoo Bay, Canada, in 2007–2008 (*Marcoux et al., 2009*). The mean rate of sightings corresponded to ~3.5 narwhals per hour of sighting effort. However, there was a substantial variation in the rate of appearance, as shown in Fig. 2a (also Fig. S1 and Video S1). A 1-hour moving average of sightings did not exceed 1.6 narwhals per hour and was null 85% of the time. Episodes of narwhal sightings could last from approximately 2.5 h to 13 h, corresponding to pods of different sizes appearing one after another (Fig. S1). The interval between consecutive sightings ranged from about a minute up to 15~28 h (Fig. 2b). It is unclear whether prolonged episodes of sightings corresponded to herds because multiple narwhal pods could be moving around the same time in opposite directions (Fig. S1).

**Fig. 2.**
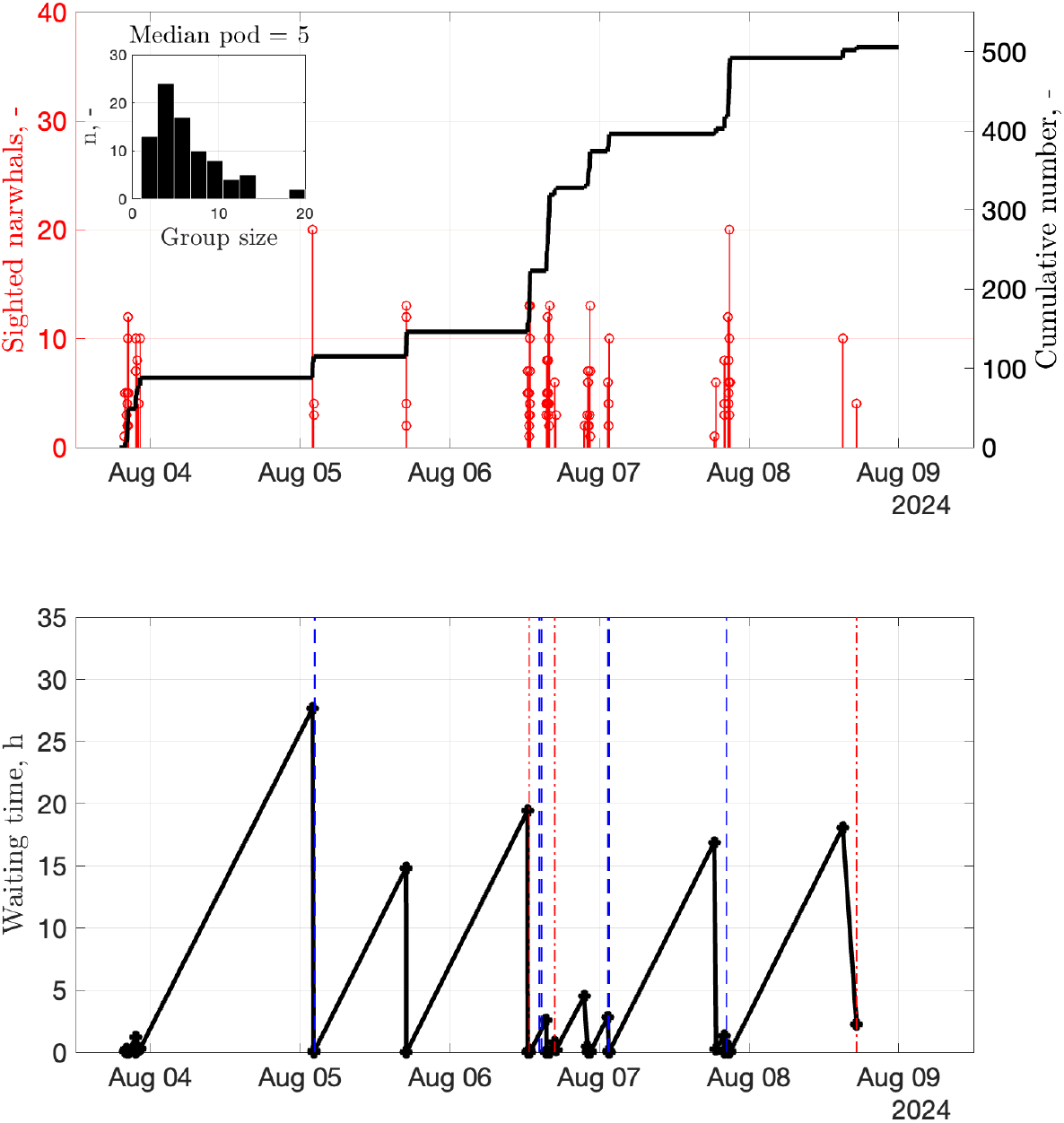
(Upper) Sighting of narwhals and cumulative number of sightings. Inset shows the distribution of pod sizes. (Lower) Waiting times between sightings. Dashed blue vertical lines indicate the timing of drone flights. Dashed red vertical lines indicate harpooning attempts. The last attempt was successful

#### Hunting attempts

The long periods spent waiting for narwhals highlight the incredible patience of Inuit hunters. Despite at least 83 encounters with narwhals during the more that six-day-long hunt, an experienced hunter made only three harpooning attempts, one of which was successful (dashed red lines in Fig. 2b); no whales were struck but lost. Such a low rate of harpooning suggests that, first, the hunt had a negligible effect on the group size, behavior, and abundance of whales in the area. Second, the hunters do not recklessly hunt narwhals, but carefully consider conditions to forecast the likelihood of hunt success and release their harpoon only when they are confident of a capture. This approach to hunting contrasts with that of modern whaling, which involves approaching and chasing whales with ships. A high attempt rate may enable hunters to catch a narwhal sooner, however, hunting failures provide narwhals with opportunities to learn about the threat posed by Inuit hunters, which may ultimately lead to a decline in the success rate over time. In the commercial whaling of sperm whales, the success rate of hunts dropped by 58% over the first few years, possibly because the whales adopted effective defensive behavior against whaling ships (*Whitehead et al., 2021*). This possibility may need to be considered for narwhals, as they have consistent migration patterns and return to Inglefield Bredning Fjord every summer, potentially with a memory of the threat from kayaks.

#### Boat front—rear difference in sightings vs. narwhal travel direction

Narwhal sightings relative to boat position are shown in Fig. 3a, with ~81% of sightings made in front of the boat (Fig. 3b). With observers communicating from all sides of the boat, including the roof, bias due to subjective perception was unlikely. However, the higher iceberg density behind the boat could have obscured the view and introduced bias. Nevertheless, the record of boat orientation suggests the boat was facing mainly east (Fig. 3a, rose diagram), with the stern towards the most ice-clad inner parts of the fjord, which might be expected to lead to fewer sightings in front of the boat (which was not the case). To aid data interpretation, sighting records were rotated based on the boat’s orientation relative to the north, aligning them with the principal orientation of Inglefield Bredning Fjord. The number of sightings and the number of sighted narwhals were associated mainly (~75%) with the eastern side of the boat, facing the head of the fjord (Fig. 3c, d). Moreover, the dominant travel direction of narwhals (Fig. 3c, rose diagram) was westward (i.e., out of the fjord; Fig. 4).

**Fig. 3.**
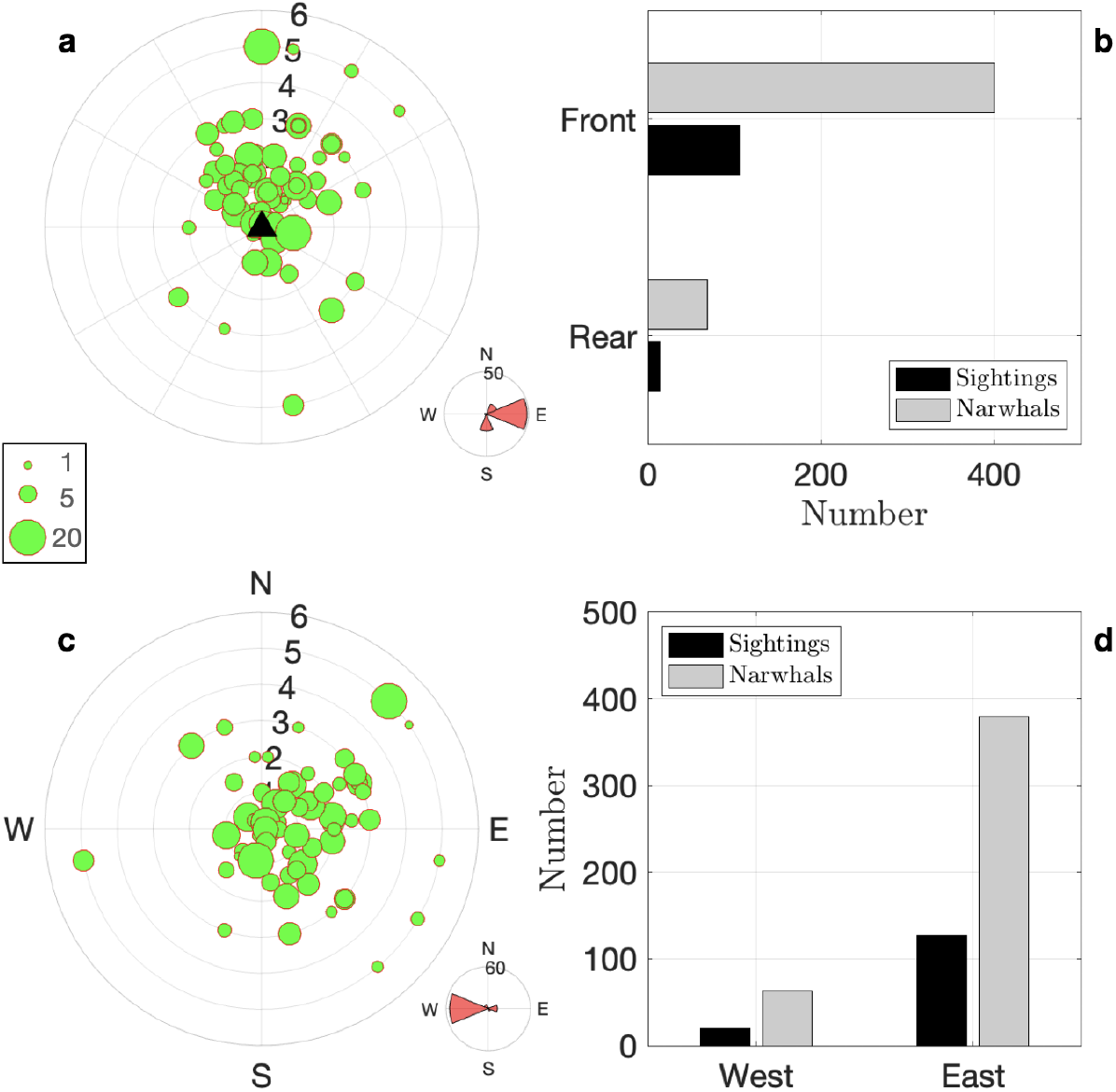
Narwhal sightings relative to the boat stern (a, b) and with rotation into cardinal directions (c, d). The circle size corresponds to the size of the group. Rose diagrams show the dominant orientation of the boat (upper) and the dominant swimming directions of narwhals (lower). The animated version is shown as Video S1

**Fig. 4.**
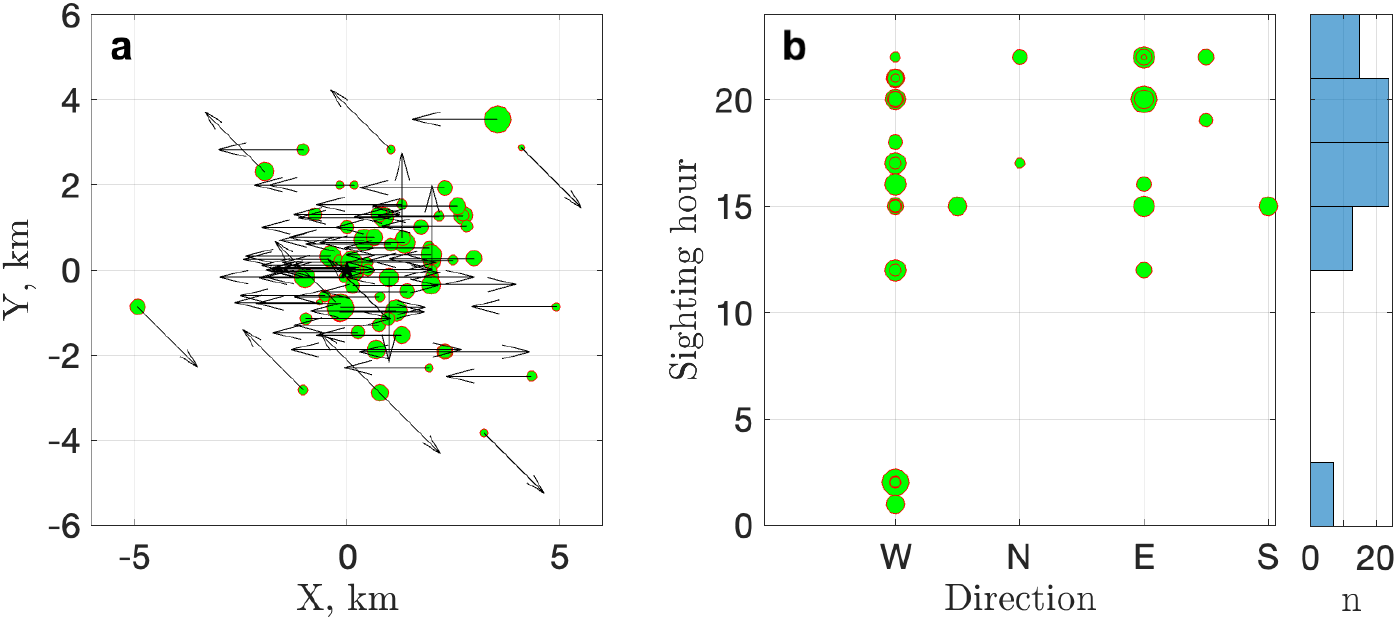
(a) Direction of narwhal-pod travel relative to the boat position (boat is at the origin; north is up). The X-axis approximately corresponds to easting; the Y-axis approximately corresponds to northing. (b) Direction of narwhal-pod travel versus the hour of sighting (with a corresponding histogram using 3-h bins). The circle size indicates the size of the group. The same data are shown vs. time, general geostrophic circulation, and tidal height in Fig. S1

The more frequently the boat was oriented eastward, the more sightings were recorded on the eastern side. Westward narwhal movement may have been influenced by the same factor that determined the boat’s orientation– namely, the near-surface outflowing current, which also guided narwhals as they passed the boat. This interpretation is consistent with previous estimates of currents undertaken in 2016 in Inglefield Bredning (*Willis et al. 2018*), showing that the general geostrophic circulation is counterclockwise (Fig. S1). Specifically, currents flow into the fjord along the southern coast and exit along the northern coast with the highest velocities (~15 cm/s) in the upper 15 m (*Willis et al. 2018*). It is possible that narwhals enter the head of the fjord south of Qeqertaq and exit the fjord along Josephine Peary Island. Earlier, *Vibe* (*1950*, p. 77) speculated that narwhals seem to migrate along the coasts into Inglefield Bredning when the tide is rising, leaving the fjord about 11 h later. However, we could not identify a clear link between our sighting data and tidal-height records (Fig. S1), possibly due to an insufficient duration of observations or a lack of tidal modulation of the current near the surface due to the strong outflow of freshwater glacier discharge that obscures the tidal signal near the surface. In fact, glacier discharge is usually stronger in the second half of the day (*Podolskiy et al., 2023*). The timing of our sightings tended to be in the second half of the day, when most narwhals traveled westward, seemingly exiting the head of the fjord (Figs. 4 and S1). The movements of other whales are known to be linked to tides and currents (e.g., *Tsujii et al., 2022*), and it is reasonable to expect that the relatively slow-swimming narwhals (*Tervo et al., 2021*) travel with geostrophic- or runoff-induced currents. Our observations may provide the first quantitative evidence of this behavior in narwhals.

Notably, the swimming direction of the narwhal is important for the hunt. When hunting narwhal, Inuit consider it and the direction of the sun. According to hunters, narwhals are sensitive to shadows, and even if hunters approach silently, narwhals can take evasive action in response to the moving shadow of the kayak. For this reason, hunters prefer a cloudy sky when the relative positions of the sun and the narwhal direction of motion are less crucial. This Indigenous knowledge suggests that visual factors such as brightness may play a role in the cognitive function of narwhals in perceiving their surrounding, in addition to sound.

### Supplementary boat/drone insights and outlook

#### Floating narwhals

The narwhal behavior of resting while floating at the surface is familiar to Inuit hunters but poorly documented. During our field observations from the boat, single or multiple narwhals were observed lying motionless on the calm water surface (n = 1 at 12:45 on 6 August 2024; n = 4 at 17:17 on 8 August 2024), with the head and back exposed and without submergence for considerable periods (Fig. 5; Video S2). In one episode, such floating behavior near the boat lasted for more than 11 min (it was interrupted when, 1.5 km away, an unsuccessful harpooning attempt of another narwhal was made). We did not aim to document this behavior from the drone due to its short battery. This immobility at the surface can last sufficiently long and enable hunters to approach the narwhals. One of four such narwhals floating motionlessly as a group was harvested during our observations. Similar floating behavior has previously been associated with rest and sleep in cetaceans and reported for multiple species (*Lyamin et al., 2008*), including bottlenose dolphins (*Tursiops truncatus*), killer whales (*Orcinus orca*), which can be motionless for more than 1 h, grey whales (*Eschrichtius robustus*), southern right whales (*Eubalaena australis*), Mediterranean fin whales (*Balaenoptera physalus*), sperm whales (*Physeter macrocephalus*), and humpback whales (*Megaptera novaeangliae*). In the North Greenlandic language (Avanersuarmiutut), such floating whales are called pugginnartoq (pronounced as ‘puhhinnarto’). As scientists know little about narwhal rest and sleep or how long narwhals need to sleep, these anecdotal accounts are useful for planning observations in the future.

**Fig. 5.**
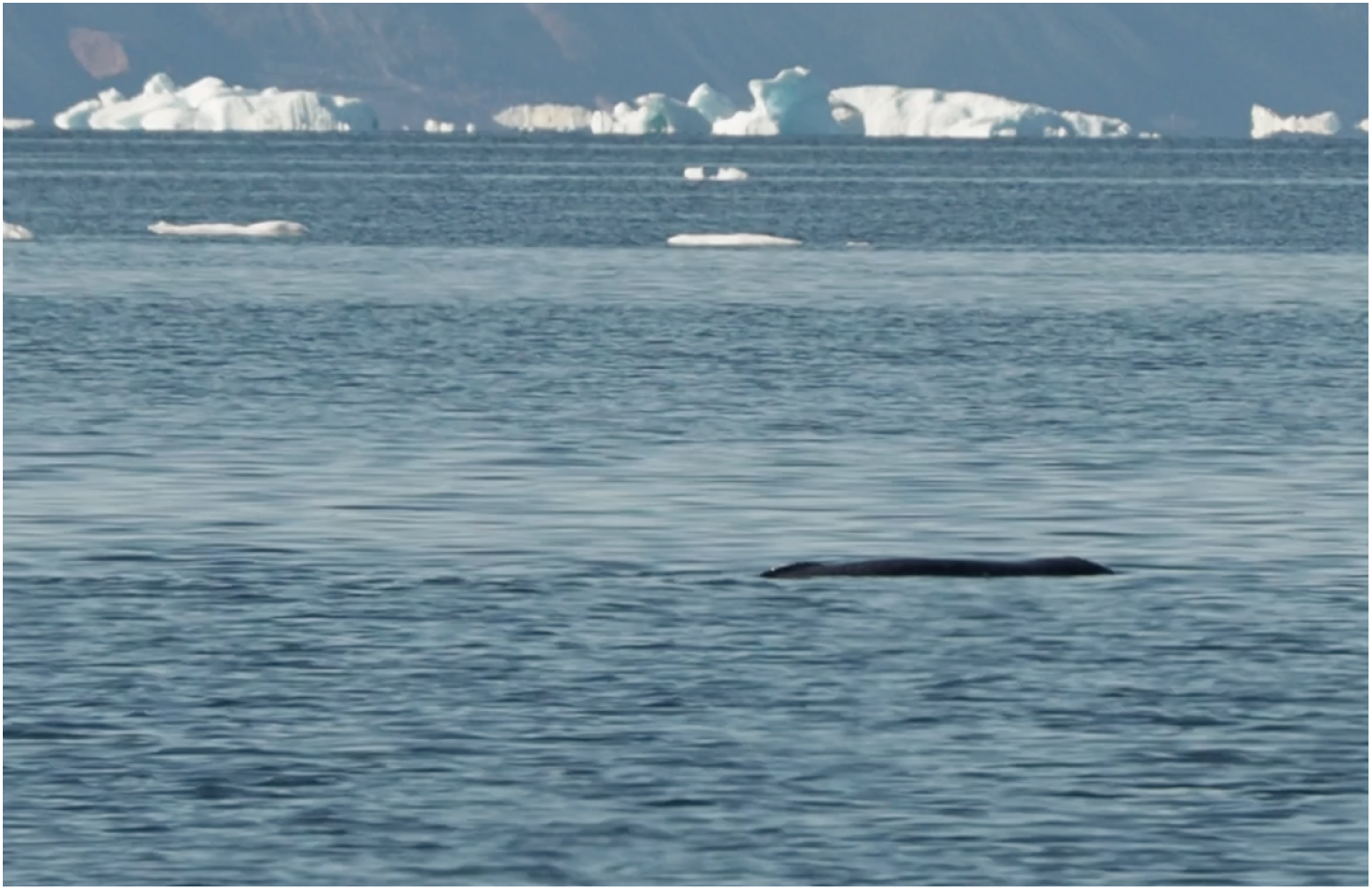
A ‘Floating’ narwhal near the boat (pugginnartoq in Greenlandic). This individual left the surface at the moment of a harpooning attempt on another narwhal at ~1.5 km distance (6 August 2024; Video S2 frame: M. Ogawa)

### Numbers of narwhals with tusks

Drone-based observations of narwhals were attempted on 5–7 August 2024 (blue lines in Fig. 2b), primarily during ‘waves’ of narwhal appearance near the boat. Fifty-eight narwhals, including two calves (~3.4%), were filmed by the drone (Fig. 6), with 37—45 having a tusk (~71% with uncertainty of ±7%). This ratio is similar to that reported by *Ogawa (2025*; 78%, 26 of 33) for narwhals harvested and sampled in 2022–2024 in the inner part of Inglefield Bredning Fjord.

**Fig. 6.**
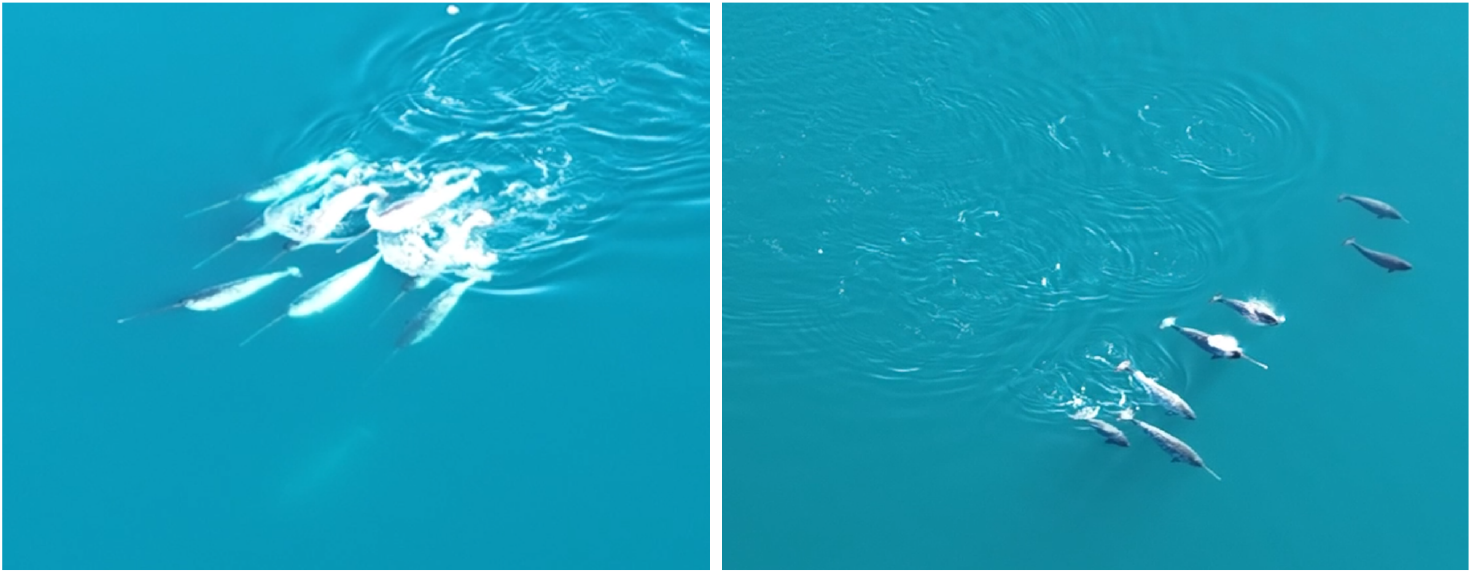
Narwhal pods comprising (left) nine mature (one is underwater at lower center) and (right) seven mixed narwhals (adults, sub-adults, and a calf) filmed from the drone on 7 August 2024 (left: altitude 61 m a.s.l., DJI_0486.MP4, frame 2336; right: altitude 70 m a.s.l., DJI_0487.MP4, frame 137). Credit: M. Ogawa

*Vibe (1950*, p. 79) reported that (i) all 20 narwhals harvested near Qeqertat in the summer of 1940 were females; (ii) of 100 narwhals caught annually at the head of Inglefield Bredning Fjord, most were females, which resurface more frequently than males; and (iii) in the outermost parts of Inglefield Bredning it was mainly traveling males that were harvested because hunters valued their tusks. Assuming all tusked narwhals are males, our data are inconsistent with Vibe’s observations near Qeqertat. Interestingly, a declining proportion of females and overrepresentation of large males was also found in East Greenland and associated with overharvest (Garde et al., 2022). However, more data are necessary to suggest any similarities.

### Tusk-to-body length ratio

Our drone footage had a dynamic angle of view intended for quick detection of narwhals, so estimation of morphological measures was not practical, although relative dimensions were considered and we attempted to extract relative differences between tusk and body lengths. We extracted frames with the most appropriate views using the Video Snapshot function in VLC Media Player (Version 3.0.21 Vetinari). For each visible tusked narwhal (n = 37), we manually estimated the length of the tusk and body in pixels using ImageJ software (version 1.54g). Snapshots from drone videos tend to not be as clear as raw images. To assess the uncertainty of this approach, we repeated measurements three times for four tusked narwhals in different frames of oblique videos (DJI_0466.MP4 and DJI_0467.MP4), yielding a mean SD of 0.01 (1%). Tusk length ranged from 11% to 56% of body length for presumably juvenile and mature narwhals, respectively. The median ratio of tusk-to-body length was 23% (Fig. 7). For comparison, five male narwhals harvested in the same area in August 2022–2024 had ratios of 17%–47% (Fig. 7; Table S1). Given that these results generally agree and considering that *Graham et al., (2020*) found a relationship between tusk length, T, and body size, B, in adult males (T=–2036.5+9.06B–0.009239B^2^), we speculate that the ratio of ~0.28 could be considered as a remotely retrievable proxy for separating adult male narwhals from juvenile (with a higher and lower ratio, respectively). Determining the age of a narwhal is a challenging task, but this cut-off could be tested with hunt data in future research.

**Fig. 7.**
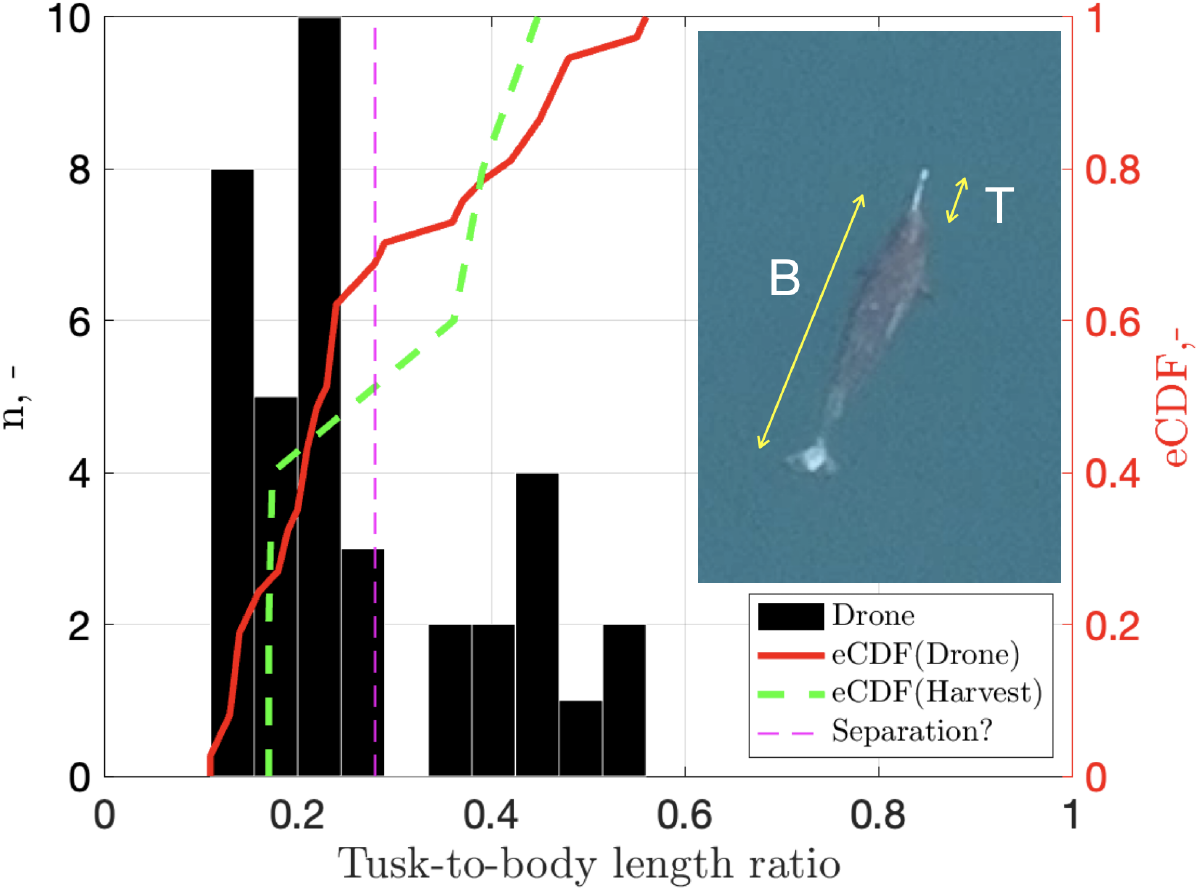
Distribution and empirical cumulative distribution function (eCDF) for the tusk (T)-to-body (B) ratio estimated from drone imagery (red) and five harvests (green) made in the same area during 2022–2024 (from Table S1). The vertical magenta line corresponds to the ratio of ~0.28, which potentially might help to separate adult male narwhals from juveniles (see the main text)

### Narwhal behavior for further investigation

Careful frame-by-frame inspection of 17 drone videos (in total, 41 min 10 s long) on a large screen (Apple Pro Display XDR 32 inches, 6K) revealed several interesting behaviors (Table S2). Narwhal groups generally maintained close spatial and temporal cohesion when surfacing and diving. Other notable behaviors included seemingly playful rolling over one another, a lack of an obvious leader in traveling and diving groups (i.e., spontaneous leadership, where even a calf may initiate a group’s dive, like in humpback-whale mothers and calves; *Ratsimbazafindranahaka et al., 2024*), gathering from different directions for joint diving, swinging of tusks, and the creating of foam (or aerated water) traces in their wake. The latter three features are detailed below, partly because they can be analyzed from static frames. We acknowledge the qualitative nature of our behavioral descriptions and present them for the reader’s consideration only as topics for further investigation with quantitative ethology methods, including time-budget analysis and AI-assisted tracking.

In three drone videos, we observed an aggregation of groups or single individuals from different directions that dived in the same location. For example, three narwhals met another two at a right angle (90°), then the three initiated a dive and were followed by the other two about 9 s later (Fig. 8). In another example, one group that dove was followed by another group at the same location after ~153 s (video DJI_0482.MP4). Finally, two individuals separated by a few hundred meters dove in the same direction with 21 s between them (video DJI_0475.MP4). Such group formation is likely an example of fission—fusion behavior in which social animals split into smaller groups and later merge back together). We believe these examples and illustrations (Fig. 8) are not only interesting by themselves, but also potentially informative for future tagging campaigns, which might reveal intermittent episodes of synchronization corresponding to the same behavior.

**Fig. 8.**
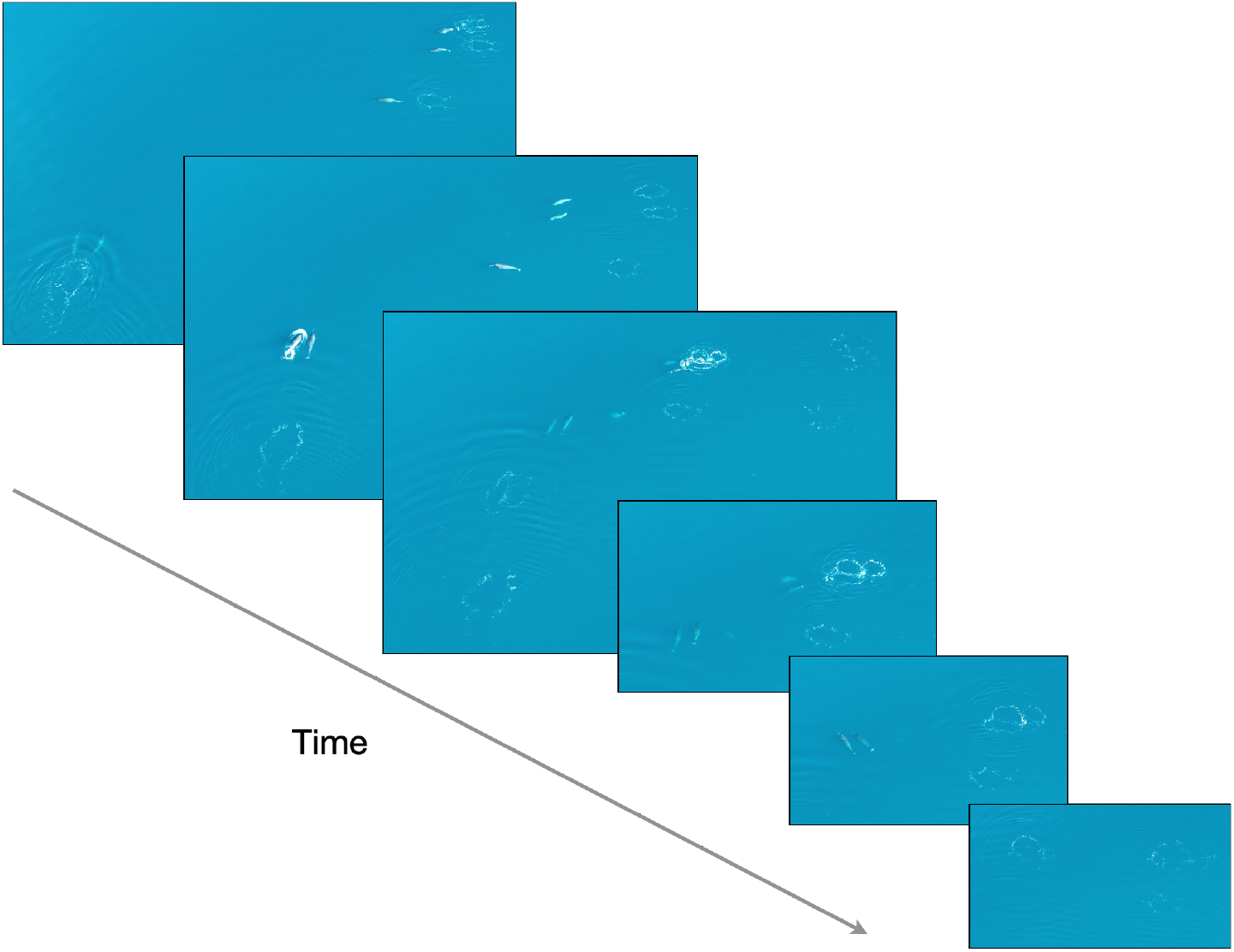
Sequence of drone images showing the meeting of two groups of narwhal to dive one after the other at the same site (DJI_0476/7.MP4; 75 m a.s.l., ~14:40 on 6 August 2024)

In several videos, we also noted that narwhals swing their tusk left and right over a range of ~30° while swimming forward (Fig. 9a). The size and quality of our videos were insufficient to quantify this. However, we observed that when a narwhal was about to meet other narwhals (e.g., Fig. 9b) or decided to follow their dive (Video DJI_0467.MP4, frame 3840), the tusk seemed to point toward them. Due to a lack of obvious points of reference, such as the tip of the tusk, tusk-less narwhals were difficult to track from the high altitudes of our videos. Nevertheless, we presume that the swinging motion likely represents echolocation scanning. Narwhals are known for some of the narrowest acoustic beams in the animal world (*Kobliz et al., 2016*), thereby requiring a wide range of motion of the upper body half to cover a larger area underwater. We also wonder if the asymmetry of the narwhal’s skull might cause a higher frequency of swings to the left than to the right. This hypothesis might be interesting to quantify in future biologging or AI-assisted posture-tracking work.

**Fig. 9.**
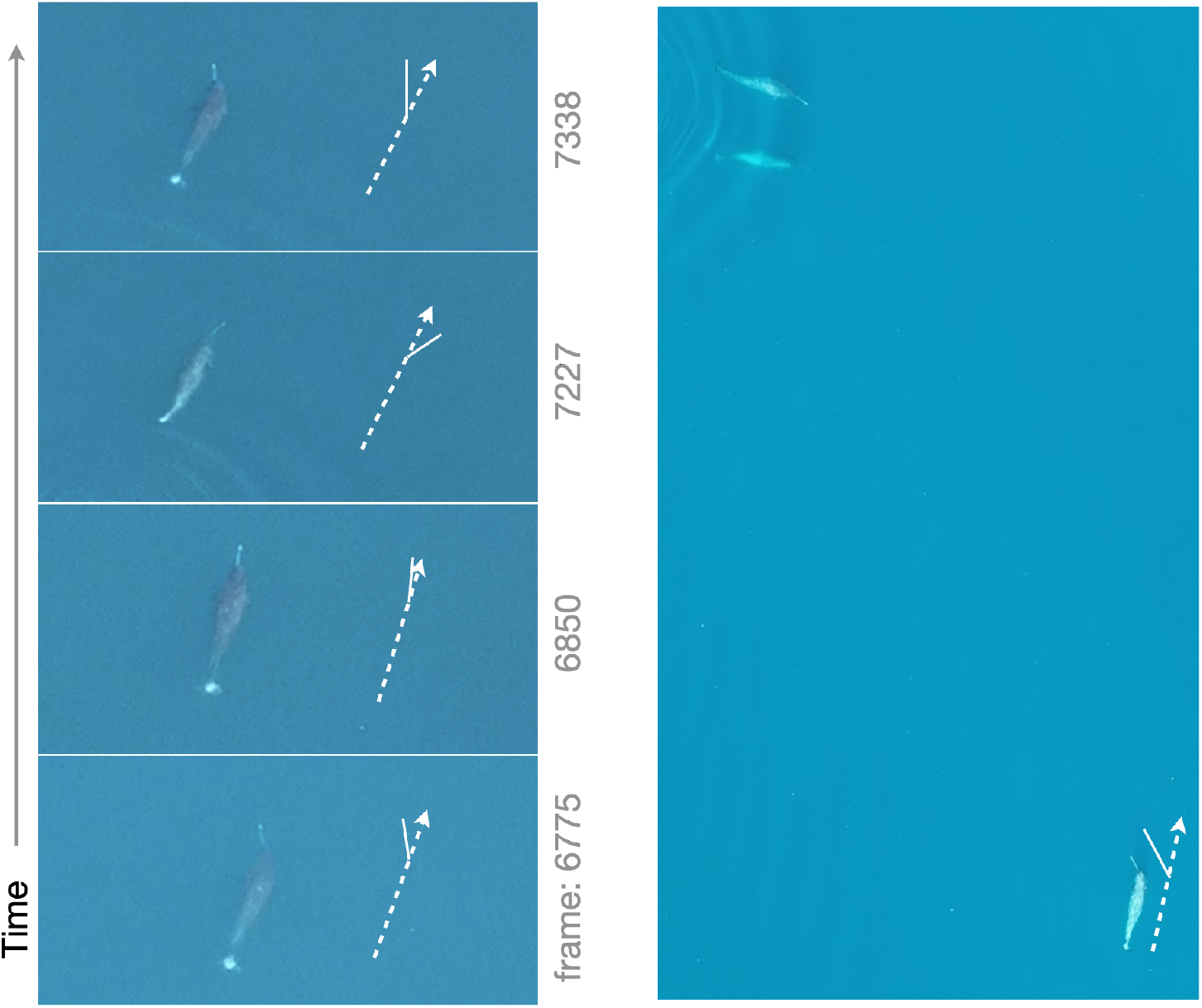
(Left) A tusk-swinging forward motion of a narwhal shown in a sequence of video frames (the first frame is from 02:23:59.244 on 5 August 2024; altitude ~67 m a.s.l., DJI_0467.MP4). (Right) A narwhal ‘looking’ toward another approaching pair (altitude ~75 m a.s.l., 14:40.40.716 on 6 August 2024, DJI_0476.MP4, frame: 7928, rotated 90° clockwise)

The drone video revealed a foam (or aerated water) trace left by traveling narwhals as a result of resurfacing and exhalation (Fig. S2) whereas the trace was barely perceptible from the water surface. Each resurfacing location became a focus of circularly expanding surface waves. These features were helpful to a drone operator in detecting narwhal presence.

## Conclusions and outlook

This field report presents several observations and lessons learned from an Inuit hunt to demonstrate that there are many different research topics that future field campaigns could strive to investigate more fully in collaboration with hunters.

We undertook boat-based observations that differed from classic marine surveys because our boat was relatively stationary. Usually, sightings are made from a moving and noisy vessel or aircraft, not allowing continuous and close observations. Inuit hunt allowed a unique perspective of narwhal behavior in their summering ground and provided a platform from which to launch a drone. Boats and drones have not previously been used for narwhal behavioral studies in this important habitat.

Over 6 days, we made 506 narwhal observations from the boat. Boat-based observations suggest that narwhals usually travel westward (outward of the head of the fjord) near the northern side of the fjord, with their occurrence being highly intermittent, tending to be between noon and midnight and lasting up to ~13 h. Sometimes they float at the surface, presumably resting or sleeping. Drone filming (58 observations) revealed a ratio of tusked-to-non-tusked narwhals of 71%±7%, with tusk-to-body ratios of 0.11–0.56 (±0.01), and documented glimpses of narwhal behavior including fission—fusion-like behavior, spontaneous leadership, tusk-swinging (presumably for scanning ahead), and a visual trace of foam at the surface, all of which are interesting topics for follow-up research. We suggest exploring these in behaviourally focused papers based on more drone videos.

Drones were useful in detecting calves and tusks, which are normally underwater when narwhals travel and therefore difficult to observe from boats. However, drone observations were limited by battery capacity and a lack of recharging facilities (a generator was not used to minimize noise). In future trials, solar panels may be used for recharging, with an abundant supply of batteries. In future work with more-stable drone videos (in terms of camera angle), we suggest using animal behavior software to create ethograms (e.g., BORIS by *Friard and Gamba, 2016*) and tracking narwhal motion with a machine learning approach (perhaps using ‘Loopy’ software; Loopbio), for analysis of speed, respiration, left—right tusk swinging, distances between individuals, and synchronization of near-surface behavior. We hope these suggestions will help to design field campaigns better.

Finally, our observations also quantitatively answer a frequently asked question: How long does it take to find a narwhal during a hunt in the ice-free season? It may take 28 h to sight a narwhal, and a week for a successful catch using traditional methods.

## Notes

### Competing Interest Statement

The authors have declared no competing interest.

### Funding

- The Sasakawa Peace Foundation: 2024-4095
- Grants-in-Aid for Scientific Research: 24K02093
- ArCS-II: JPMXD1420318865

