## Supplemental Materials for "Inuit Hunt as a Platform for Observing Narwhals (*Monodon monoceros*) in Inglefield Bredning (Kangerlussuaq), Greenland"

**Supplementary Information (SI)**

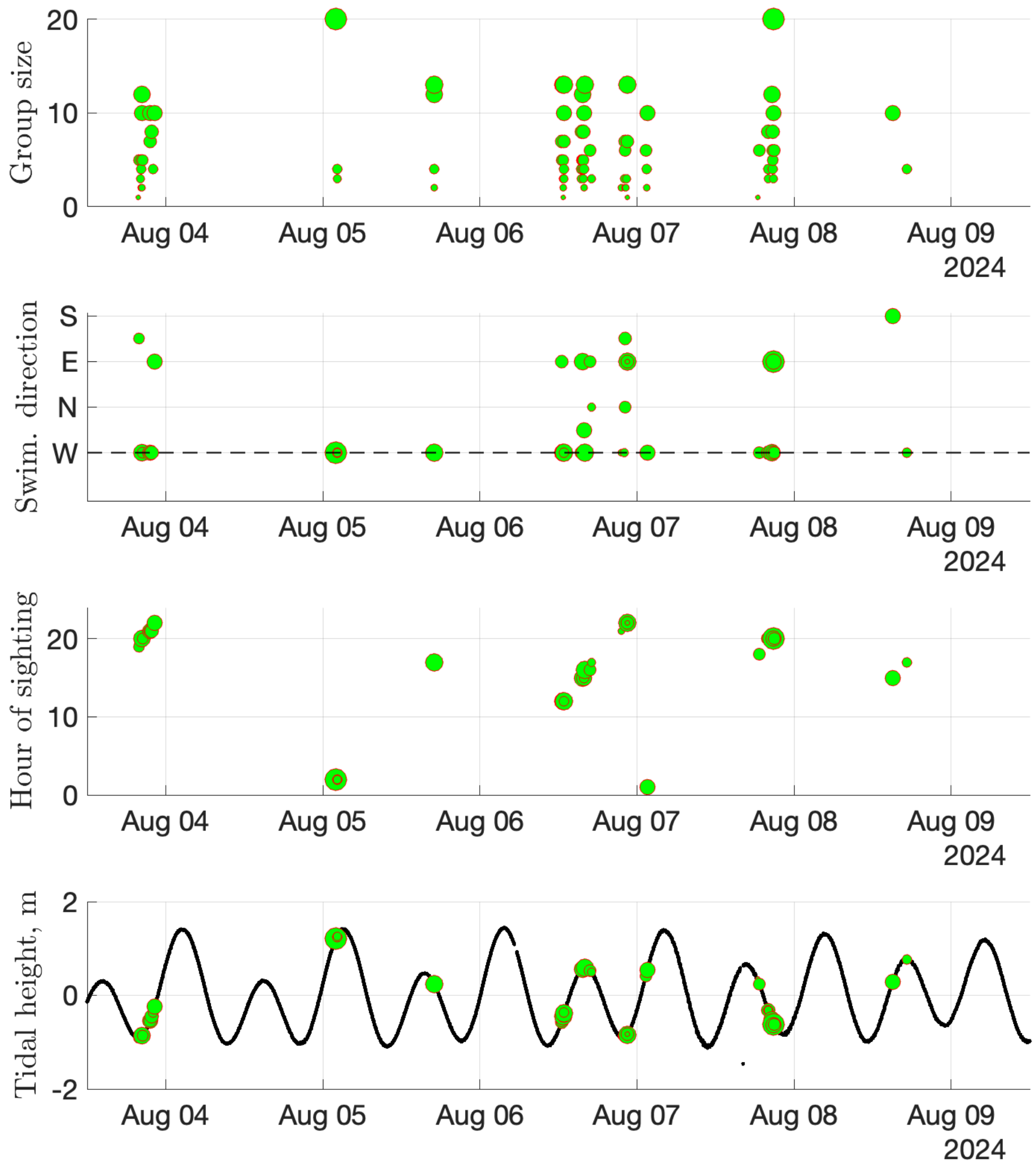


***Fig. S1.*** *Timing of narwhal pod sightings compared to their pod size, direction of travel (with the direction of the general geostrophic circulation shown by a dashed line), hour of pod sighting, and tidal height at Thule—Pituffik station (data: https://www.ioc-sealevelmonitoring.org/station.php). Previous work confirmed the local tide at the site of the field expedition is in phase with that at Thule (Minowa et al., 2019).*

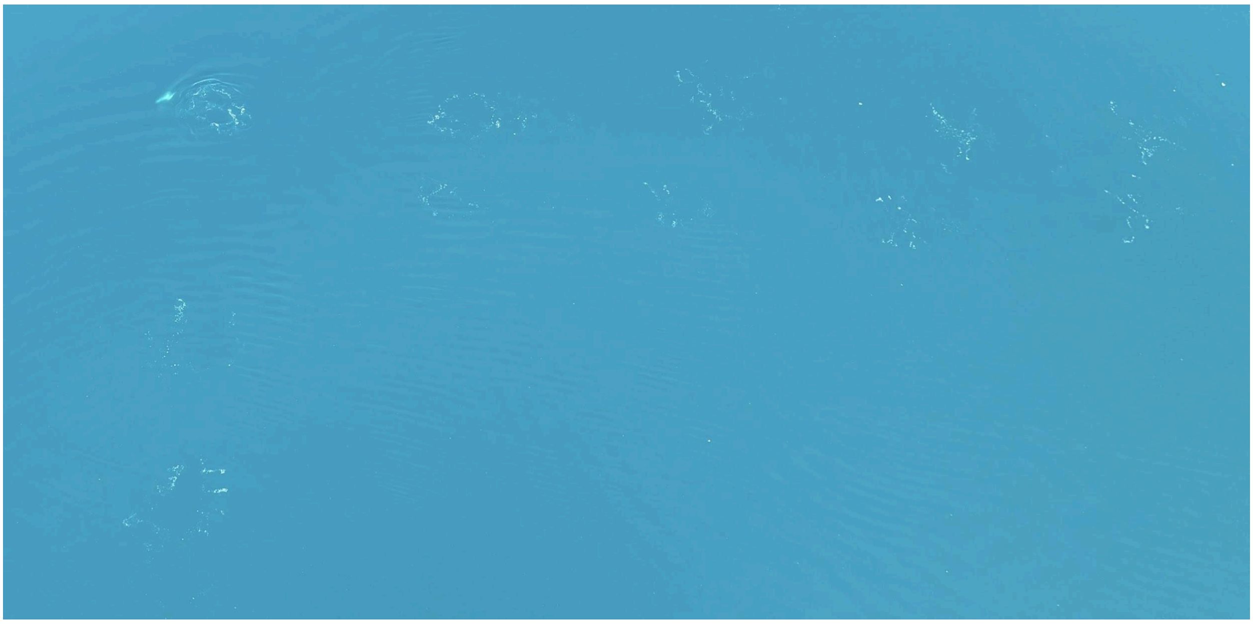

***Fig. S2.*** *Foam traces from two pairs of narwhals meeting at a right angle and diving in the area at upper left in the image (altitude ~75 m a.s.l., DJI_0477.MP4). Credit: M. Ogawa.*

**Table S1.** Body size and tusk length of male narwhals harvested in Inglefield Bredning (Kangerlussuaq) in summers 2022–2024.

| Catch year | Body size, cm | Tusk length, cm |
|---|---|---|
| 2022 | 460 | 180 |
| 2023 | 453 | 203 |
| 2023 | 418 | 151 |
| 2023 | 299 | 52 |
| 2024 | 370 | 63 |

**Table S2.** Drone-video file notes (data accessibility statement shows links to the videos).

| **Video file (a few if a sequence) with time-stamp** | **Narwhals, n (with tusk)** | **Remarks** |
|---|---|---|
| DJI_0466.MP4 (>02:21, 5 August 2024) | 4 (4) | Three travelling narwhals "scan" by swinging tusks and make a shallow dive; later they resurface with a fourth individual. |
| DJI_0467.MP4, DJI_0468.MP4 (>02:24, 5 August 2024) | 4 (4) The same group as above (thus not included in total counts), but one of the narwhals differs from the original group 1(1). | Three narwhals dive (one rolles other another, frame 2961), the fourth narwhal looked in their direction, took a breath and followed (12 sec later); all seemed to swing tusks. Later another narwhal appears, seems to search, and dives. Several little auks nearby are also diving. |
| DJI_0472.MP4 (>14:18, 6 August 2024) | 1 (?) | Narwhal dives; afterward, a clear foam trace remains (for at least 3 minutes). |
| DJI_0474.MP4, DJI_0475.MP4 (>14:23, 6 August 2024) | 4 (2) - including a calf; we can see another remote group of ~6 (tusks were unclear), and later one non-tusked narwhal. | Two pods travel between icebergs. One pod is too far to clearly see. Three narwhals from the closer pod dive (calf leads, DJI_0474.MP4, frame: 6462), and one tusked narwhal remains at the surface. A few hundred meters away, there is another non-tusked narwhal at the surface approaching. The tusked narwhal dives. About 21 seconds later, the non-tusked narwhal seems to dive in the same direction. |
| DJI_0476.MP4, DJI_0477.MP4 (>14:41, 6 August 2024) | 1 (?)<br>3 (2)<br>2 (2) | Single narwhal resurfaces (at 48 sec). Later, a group of three meets a pair at a right angle and they all dive steeply with a trace of surface bubbles. Appears to be fusion behavior. |
| DJI_0478.MP4 (>14:46, 6 August 2024) | 5 (4) - the same group as above observed shortly afterwards (thus not included in total counts) | Group dive, led by one narwhal without a tusk, who takes an extra breath whereas others do not. |
| DJI_0481.MP4,<br>DJI_0482.MP4,<br>DJI_0483.MP4<br>(>01:24, 7 August 2024) | 3 (2)<br>5 (3) - view another remote group<br>5 (5) | Three narwhals travel and dive, one with tusk leads. This dive is followed by two closely moving pods at a distance of a few hundred meters (another far at the back; closer to boat). One pod |

| | | |
|---|---|---|
| | | dives first (70 s after the first narwhal), followed by the second pod (83 s after the first pod). |
| DJI_0484.MP4 (>01:33, 7 August 2024) | 4 (3) | Spontaneous leadership; “scan” by swinging tusks and rotating, then dived. |
| DJI_0486.MP4, DJI_0487.MP4 (>20:24, 7 August 2024) | 9 (8+1?)<br>7 (3) - including a calf | A second group seen at the back in the beginning, group of nine dive. Frame >8000, another group of seven, dives together (seems like escape). After that, see six and a calf staying next to white mother. |
| DJI_0488.MP4 (>20:30, 7 August 2024) | 1(1) | single travelling narwhal |
| DJI_0489.MP4 (>20:31, 7 August 2024) | 1(1) | single travelling narwhal |
| Total = | 58 (min & max = 37 & 45) | 71±7% of narwhals had tusks |

**Dataset S1** (separate file). Sighting data (3—8 August 2024).

### Multimedia files

**Video S1** (separate file). Animated time variation of narwhal sightings relative to the boat (3—8 August 2024).

**Video S2** (separate file). “Floating” narwhal near the boat (6 August 2024; Credit: M. Ogawa).
